# Glycerophosphodiesterase GDE2 affects pancreas differentiation in zebrafish

**DOI:** 10.1101/108779

**Authors:** Michiel van Veen, Jason van Pelt, Laurie A. Mans, Wouter H. Moolenaar, Anna-Pavlina G. Haramis

## Abstract

Notch signaling plays an essential role in the proliferation, differentiation and cell fate determination of various tissues, including the developing pancreas. One regulator of the Notch pathway is GDE2 (or GDPD5), a transmembrane ecto-phosphodiesterase that cleaves GPI-anchored proteins at the plasma membrane, including a Notch ligand regulator. Here we report that Gde2 knockdown in zebrafish embryos leads to developmental defects, particularly, impaired motility and reduced pancreas differentiation, as shown by decreased expression of insulin and other pancreatic markers. Exogenous expression of human GDE2, but not catalytically dead GDE2, similarly leads to developmental defects. These data reveal functional conservation between zebrafish and human GDE2, and suggest that strict regulation of GDE2 expression and catalytic activity is critical for correct embryonic patterning. In particular, our data uncover a role for GDE2 in regulating pancreas differentiation.

## Introduction

A better understanding of pancreatic homeostasis is essential to identify new ways to increase insulin-producing beta cell number and to enhance their function, in order to tackle diabetes, a rising epidemic.

The Notch cell-to-cell signaling axis plays an important role in proliferation, differentiation and cell-fate decisions in the developing pancreas and other tissues (Ninov et al., 2012;Tehrani et al., 2011). Notch receptor signaling is activated by transmembrane Notch ligands in adjacent cells (Wang et al., 2011). Following ligand binding, Notch is cleaved by the metalloprotease TACE (factor-a-converting enzyme) (Wang et al., 2011). Truncated Notch is in turn cleaved by the a secretase complex, leading to intracellular release of NICD (Notch intracellular domain) (Wang et al., 2011) and activation of Notch target genes (Wang et al., 2011).

Different levels of Notch can elicit distinct fates (Ninov et al., 2012;Tehrani et al., 2011). In general, Notch signaling maintains progenitor cells in an undifferentiated proliferative state and regulates timing of differentiation, not only in the embryonic pancreas but also in developing motor neurons (Sabharwal et al., 2011;Zecchin et al., 2007). Loss of Notch signaling in mice (Apelqvist et al., 1999) and zebrafish (Esni et al., 2004;Zecchin et al., 2007) results in aberrant/excessive differentiation of pancreatic progenitors to endocrine cells, at the expense of the later-appearing exocrine cells (Tehrani et al., 2011). Modulation of Notch signaling may thus affect the balance between proliferation and differentiation of pancreatic progenitor cells.

A recently identified modulator of Notch activity is GDE2 (aka GDPD5), a multipass membrane glycoprotein with an extracellular glycerophosphodiesterase (GDPD) domain. The GDE2 catalytic domain cleaves a subset of glycosylphosphatidylinositol (GPI)-anchored proteins, including a Notch ligand regulator (RECK) (Park et al., 2013) and certain heparan sulfate proteoglycans, the glypicans (Cave et al., 2017; Matas-Rico et al., 2016a). Trough RECK cleavage, GDE2 sheds and inactivates a Notch ligand (Delta-like 1) in the developing spinal cord, leading to Notch inactivation in adjacent progenitor cells (Park et al., 2013;Sabharwal et al., 2011). Thus, GDE2 promotes motor neuron differentiation by downregulating Notch signaling during spinal cord development. Mice lacking GDE2 exhibit selective losses of limb-innervating motor neuron pools leading to neurodegeneration (Cave et al., 2017;Sabharwal et al., 2011). In addition, GDE2 can promote neuronal differentiation in a cell-autonomous manner, namely through GPI-anchor cleavage of glypican-6, which correlates with improved clinical outcome in neuroblastoma (Matas-Rico et al. 2016a(Matas-Rico et al., 2016b). However, a possible role for GDE2 in regulating the fate of non-neuronal cell types has not been investigated to date.

Here, we examine the role of Gde2 during embryonic development, focusing on pancreas differentiation in zebrafish. Zebrafish is an excellent model to conduct developmental studies owing to the availability of large numbers of readily accessible, transparent embryos. Importantly, mammalian and zebrafish pancreas share many morphological and physiological similarities (Tehrani et al., 2011), while important genes in mammalian islet development are functionally conserved in zebrafish (Tehrani et al., 2011). The conservation of signaling pathways and mechanisms of pancreas development suggests that results found in zebrafish will advance our understanding of pancreas development in humans. We show that Gdpd5a knockdown affects the differentiation of specific endocrine and exocrine progenitors. Furthermore, exogenous expression of human GDE2, but not of catalytically dead mutants leads to developmental abnormalities. These results highlight the functional conservation of GDE2 and suggest that tight regulation of Gdpd5a levels and catalytic activity are important for correct embryonic patterning.

## Results

### Analysis of zebrafish Gde2

GDE2, encoded by *GDPD5,* is a six-transmembrane-domain protein (GDE2) with an extracellular phosphodiesterase domain and intracellular N- and C-termini (Fig. 1A). GDE2 catalytic activity has long been elusive, but recent studies have shown that GDE2 cleaves GPI-anchored proteins at the plasma membrane to drive neuronal differentiation and survival, in either a cell-autonomous or non-cell-autonomous manner (Cave et al., 2017;Matas-Rico et al., 2016a;Park et al., 2013) (Fig. 1A).

**Figure 1.**
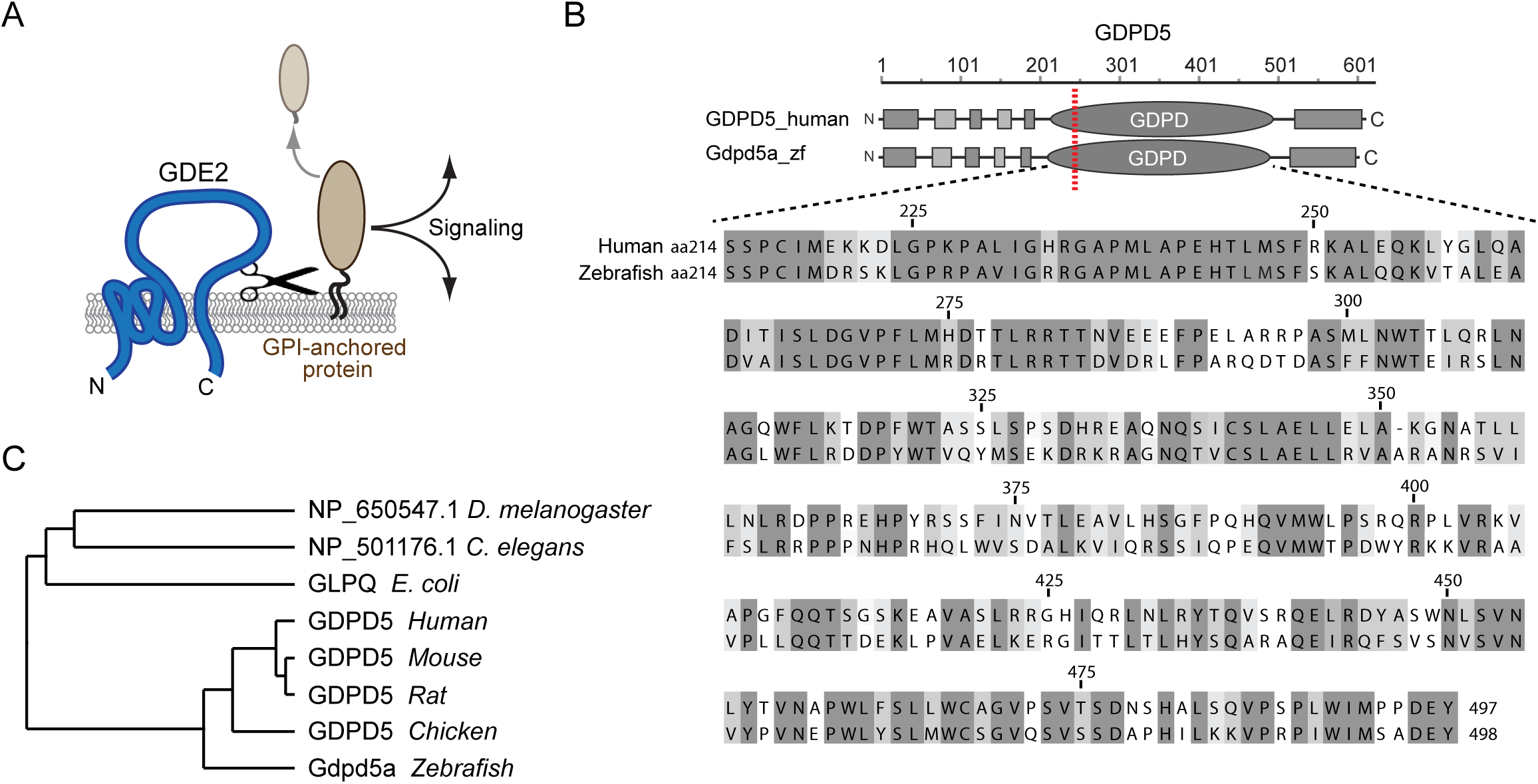
Sequence analysis of zebrafish Gdpd5a. **(A)** GDE2 cleaves GPI anchored proteins at the plasma membrane resulting in their shedding with consequent activation of signalling pathways, either in the same cell or in adjacent cells. (**B)** Amino acid sequence comparison between human GDPD5 and zebrafish Gdpd5a. The red line depicts key catalytic residue H^233^ in GDE2 (Matas-Rico et al., 2016a). Sequence comparison of the ecto-domain using Clustal Omega, shows identical residues in dark grey and similar residues in light grey. The catalytic domains share 52% sequence identity and 82% similarity. **(C)** Phylogenetic analysis showing that zGdpd5a is closely related to mammalian GDPD5. Protein sequences used: hGDPD5 (NP_110419.5), mGDPD5 (NP_958740.2), rGDPD5 (XP_006223480.1), cGDPD5 (NP_001032348.1), zGDPD5 (XP_005157799.1), *D. melanogaster* GDPD5 (NP_650547), *C. elegans* GDPD5 (NP_501176), *E. coli* GDPD5 (NP_416742.1).

Human *GDPD5* has two homologues in the zebrafish genome, *Gdpd5a* and *Gdpd5b. Gdpd5a* (glycerophosphodiester phosphodiesterase domain containing 5a) gives rise to three alternatively spliced isoforms; Isoforms *gdpd5a-*001 and *gdpd5a-*002 are included within the long isoform *Gdpd5a-*201 which encodes for 595-aa protein. Phylogenetic analysis shows that the long isoform of zebrafish *Gdpd5a* is closely related to human, rodent and chicken *GDPD5/Gdpd5* (Fig. 1B). The predicted catalytic domains of human GDE2 and zebrafish Gdpd5a share 53% identity and 81% similarity (Fig. 1C). *Gdpd5b* (ENSDARG00000076962) encodes a 566-aa protein that is characterized by a relatively short, truncated C-terminal tail (SFig. 1).

### Gdpd5a depletion leads to developmental defects

To investigate the function of Gdpd5 during zebrafish embryonic development, we knocked down its function by using antisense morpholino oligonucleotides (MO) against *gdpd5a-*201 mRNA. The MO targets the 5′UTR of the long isoform, *Gdpd5a-*201, and blocks translation of the message (Nasevicius et al., 2000). Since the other isoforms are included in *Gdpd5a-*201, we reasoned that this MO would block translation of all isoforms.

Zebrafish embryos were injected with various amounts of *gdpd5a* MO at the 1-2 cell stage and development of the injected embryos was followed up to 5 days post-fertilization (dpf). Injections with low amounts (< 8,4 ng/ul) MO did not result in morphological abnormalities. Following injection of 1 mM *gdpd5a*MO, injected embryos exhibited developmental defects. Specifically, the majority of *gdpd5a*MO-injected embryos exhibited heart abnormalities accompanied by heart oedema, as well as a shorter and bent body axis at 3 dpf (Fig. 2). Furthermore, while at 3 dpf almost all non-injected controls have hatched from their chorions, the majority of *gdpd5a*MO-injected embryos were still in their chorions, indicating inability to hatch (SFig. 2). At 5 dpf, very few of the *gdpd5a*MO-injected embryos had hatched. The Gdpd5a-depleted larvae that hatched (either by themselves or following manual dechorionation) showed severe motility defects, either swimming inability or trembling (data not shown), consistent with *Gdpd5* knockout causing behavioral motor deficits in mice (Cave et al., 2017).

**Figure 2.**
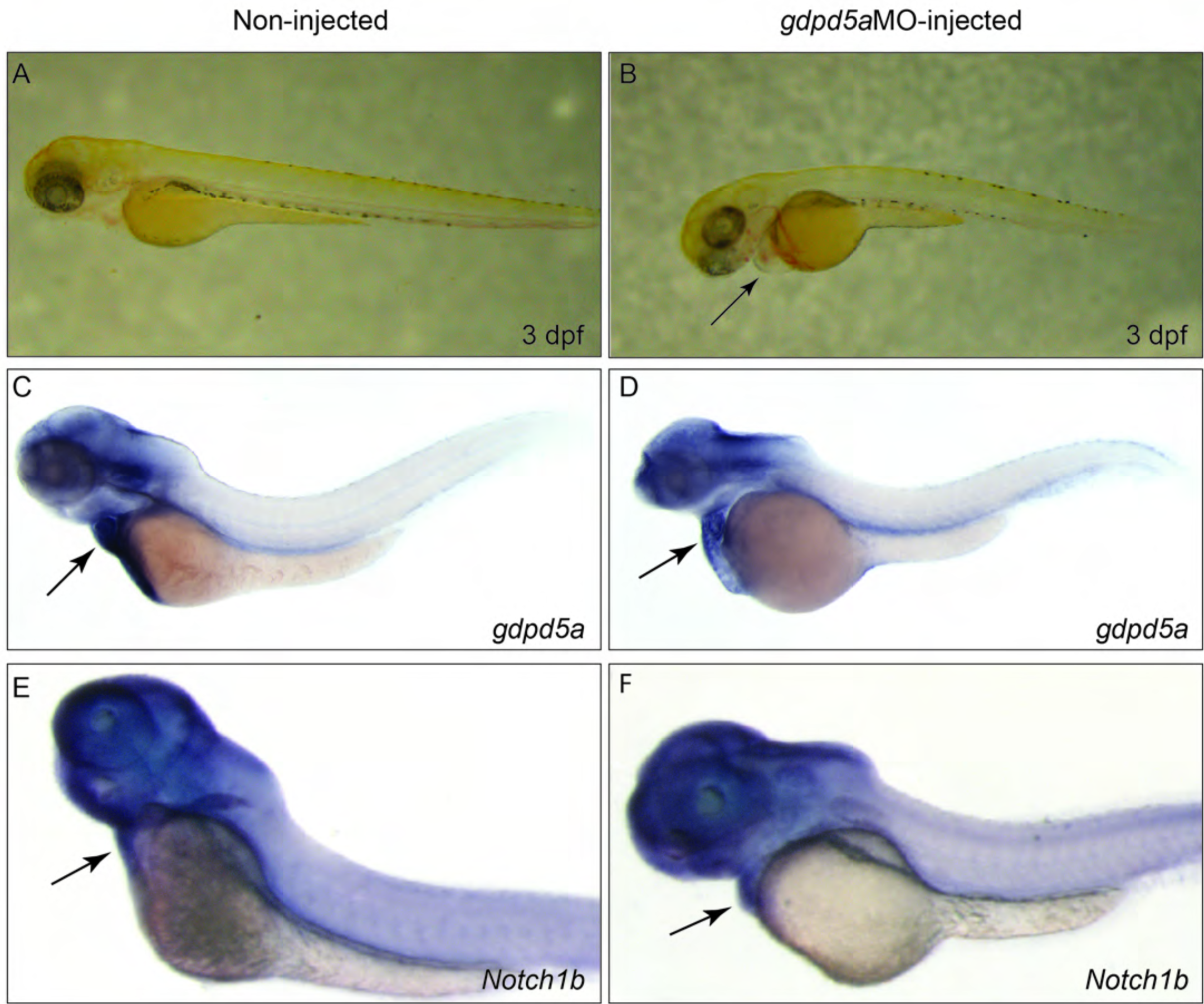
Gdpd5a depletion causes developmental defects. **(A)** Un-injected embryo at 3 dpf. Arrow indicates looped heart. **(B)** Representative example of a *gdpd5a*MO-injected embryo at 3 dpf. *Gdpd5a*MO-injected embryos show a bent and shorter body axis, heart oedemas (arrow) and unlooped heart. Whole-mount *in situ* hybridization (WISH) for expression of *gdpd5a* and *Notch1b* in non-injected wt **(C, E)** and gdpd5aMO-injected embryos **(D, F)** at 3 dpf. Lateral views. **(C)** wt embryo at 3 dpf stained for *gdpd5a. Gdpd5a* expression is detected in the brain, strongly in the heart (arrow) as well as the intestine. **(D)** Gdpd5MO-injected embryo at 3 dpf stained for *gdpd5a. Gdpd5a* is expressed at comparable levels as in non-injected controls, except for reduced expression in the heart (arrow). **(E)** wt embryo at 3 dpf stained for *Notch1b.* Strong expression in the head and heart (arrow) is detected **(F)** Gdpd5MO-injected embryo at 3 dpf stained for *Notch1b.* The expression pattern of *Notch1b* is similar to that of the non-injected controls.

### Expression studies

We next assayed *gdpd5a* and *Notch1b* expression in *gdpd5a*MO-injected embryos at 3 dpf. We cloned part of the *gdpd5a* coding region, made an antisense riboprobe against *Gdpd5a-*201 and analysed expression by *in situ* hybridization. In embryos at 3 dpf, *gdpd5a* is found expressed in the brain and intestine, and at high levels in the heart and associated blood vessels (Fig. 2C). This localization pattern suggests a role for *gdpd5a* during differentiation of these organs. The MO against *gdpd5a* is targeted to the 5′ UTR including the ATG start codon and should block translation and hence it is not expected to influence mRNA levels. Nonetheless, we detected reduced expression of *gdpd5a* mRNA in gdpd5aMO-injected embryos, particularly in the heart (Fig. 2D), suggesting that the MO may affect the stability of the *gdpd5a* mRNA. We also assayed *Notch1b* expression in *gdpd5a* morphants. *Notch1b* is expressed highly in brain and heart of 3 dpf un-injected embryos (Fig. 2E). *Notch1b* expression was comparable in *gdpd5a* morphants (Fig. 2F).

### Pancreas differentiation is affected upon Gdpd5a depletion

Given the role of Notch1b signaling in pancreas differentiation and the purported role of Gdpd5 in modulating Notch activity, we examined how Gdpd5a depletion affects pancreas differentiation in zebrafish embryos. The pancreas in zebrafish develops from the posterior foregut endoderm; the dorsal bud which produces endocrine cells emerges after 24 hpf, and the ventral bud emerges at 32 hpf and produces mostly exocrine and some endocrine cells (Field et al., 2003). At around 72 hpf the pancreas is fully developed (Tehrani et al., 2011). At 14 hrs post-fertilization (hpf) the pancreatic progenitor cells express different levels of pdx1 (Pancreatic and duodenal homeobox 1) (Tehrani et al., 2011). Cells expressing high levels of *pdx1* give rise to the endocrine cells while cells with lower *pdx1* expression give rise to exocrine and intestinal cells. The first expression of insulin begins at 15 hpf when the insulin-producing cells begin to migrate towards the midline (Tehrani and Lin, 2011). In the mature pancreas, insulin is expressed exclusively in the beta cells.

We performed *in situ* hybridizations using markers of endocrine and exocrine pancreas components in wt and *gdpd5a* morphants at 3 dpf. In gdpd5aMO-injected embryos insulin (*ins*) expression was greatly reduced (Fig. 3A). *Pdx1* was expressed in the head, intestine and endocrine pancreas at 3 dpf (Fig. 3C and SFig. 3A). No differences in *pdx1* expression were observed between non-injected and gdpd5aMO-injected embryos at 3 dpf (Fig. 3D and SFig. 3B). We therefore conclude that the defect in insulin expression is specific.

**Figure 3.**
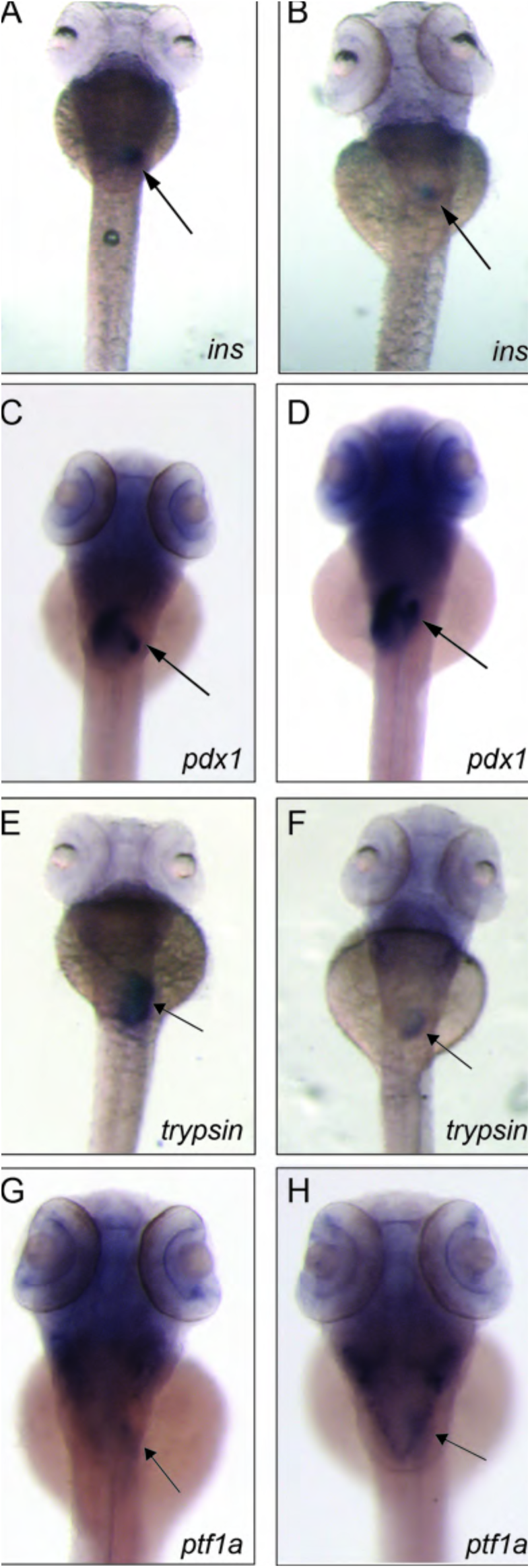
Expression of *insulin* and *trypsin,* but not *pdxl* or *ptfla,* is reduced upon Gdpd5a knockdown. WISH for expression of the endocrine pancreas markers, *insulin* (*ins*) and *pancreatic and duodenal homeobox 1* (*pdx1*) and the exocrine pancreas markers, *trypsin* and *ptf1a* in non-injected wt (A, C, E, G) and gdpd5aMO-injected embryos at 3 dpf (B, D, F, H). Dorsal views. **(A)** wt embryo at 3 dpf stained for *insulin.* Strong expression is detected in the endocrine pancreas (arrow). **(B)** Gdpd5MO-injected embryo at 3 dpf. *ins* expression is dramatically reduced. **(C)** wt embryo at 3 dpf stained for *pdx1. Pdx1* is expressed in the intestine and endocrine pancreas (arrow). **(D)** *Pdx1* expression in the intestine and endocrine pancreas is very similar in *gdpd5*MO-injected embryos at 3 dpf. **(E)** wt embryo at 3 dpf stained for *trypsin.* Strong expression is detected in the exocrine pancreas (arrow). **(F)** *Gdpd5*MO-injected embryo at 3 dpf. *Trypsin* expression is dramatically reduced. **(G)** wt embryo at 3 dpf stained for *ptf1a. Ptf1a* expression is detected in the brain and the exocrine pancreas (arrow). **(H)** *Gdpd5*MO-injected embryo at 3 dpf. *Ptf1a* is expressed at comparable levels in the brain and exocrine pancreas (arrow) as in non-injected controls.

We next examined exocrine pancreas development in Gdpd5a-depleted embryos. *Trypsin* and *pancreas-specific transcription factor 1a* (*ptf1a*) mark the exocrine pancreas. As shown in Fig. 3E, *trypsin* expression was dramatically reduced in gdpd5aMO-injected embryos compared to un-injected controls. *Ptf1a* was strongly expressed in the head and the exocrine pancreas in un-injected controls at 3 dpf (Fig. 3G and SFig. 3C). The *ptf1a* expression pattern was not altered in *gdpd5aMO-* injected embryos (Fig 3H and SFig. 2D).

Since Gdpd5a-depleted embryos exhibit mobility defects and impaired pancreas differentiation, while *Gdpd5* knockout in mice leads to motor neuron defects (Sabharwal et al., 2011), we examined expression of *islet1* in *gdpd5a* morphants. The transcription factor Islet1 is involved in both motor neuron differentiation and pancreas differentiation in mice and zebrafish (Appel et al., 1995) (Dasen et al., 2009). We found *islet1* expression in the brain, spinal motor neurons and the pancreas in wt embryos at 3 dpf (Fig. 4A, C). In *gdpd5a* morphants, *islet1* was expressed at high levels in the brain and pancreas (Fig. 4 B, D). However, in the ventral spinal cord, *islet1* expression was reduced upon *gdpd5a* knockdown (Fig. 4B).

**Figure 4.**
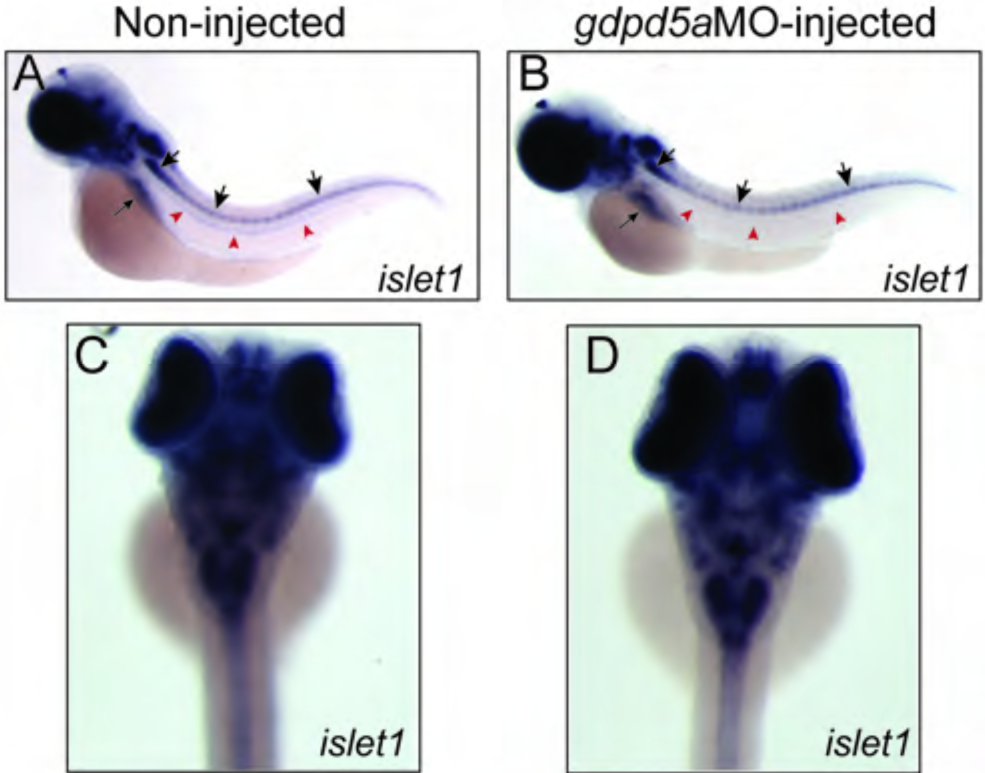
*Islet1* expression upon Gdpd5 knockdown. WISH hybridization for *islet1* expression on non-injected **(A, C)**, or gdpd5MO-injected embryos **(B, D)** at 3 dpf. **(A, B)** Lateral views, anterior to the left. **(A)** wt embryo at 3 dpf, *Islet1* is expressed in the brain, spinal cord (arrowheads) and the pancreas (arrow). **(B)** *Islet1* is expressed at similar levels in *gdpd5*MO-injected embryo. **(C, D)** Dorsal views of the head region. **(C)** wt embryo at 3 dpf. *Islet1* is expressed at distinct domains in the brain. (**D)** *Gdpd5*MO-injected embryos at 3 dpf show a staining pattern similar to that in non-injected controls.

### Expression of human GDE2 in zebrafish leads to developmental defects

To investigate the functional conservation between human and zebrafish GDE2, we injected mRNA encoding human GDE2 tagged with HA in zebrafish embryos at the one-cell stage. Expression of human GDE2-HA mRNA led to developmental abnormalities in a dose-dependent manner (Fig. 5B, C). Specifically, hGDE2 mRNA-injected larvae at 4 dpf showed malformations in the body axis and defects in the vasculature (Fig. 5C). Higher doses resulted in more severe disruption of the vasculature, blood accumulation and heart oedemas (Fig. 5C). Importantly, expression of catalytically dead human GDE2 (GDE2^His233Ala^ or GDE2^His233Ala/His275Ala^) (Matas-Rico et al., 2016a) did not induce morphological abnormalities (Fig. 5D). We verified that wt hGDE2 and GDE2^His233Ala^ and GDE2^His233Ala/His275Ala^ are expressed in zebrafish at comparable levels (Fig. 5E). These data indicate that the malformations observed depend on the catalytic activity of GDE2, and they confirm functional conservation between human and zebrafish GDE2.

**Figure 5.**
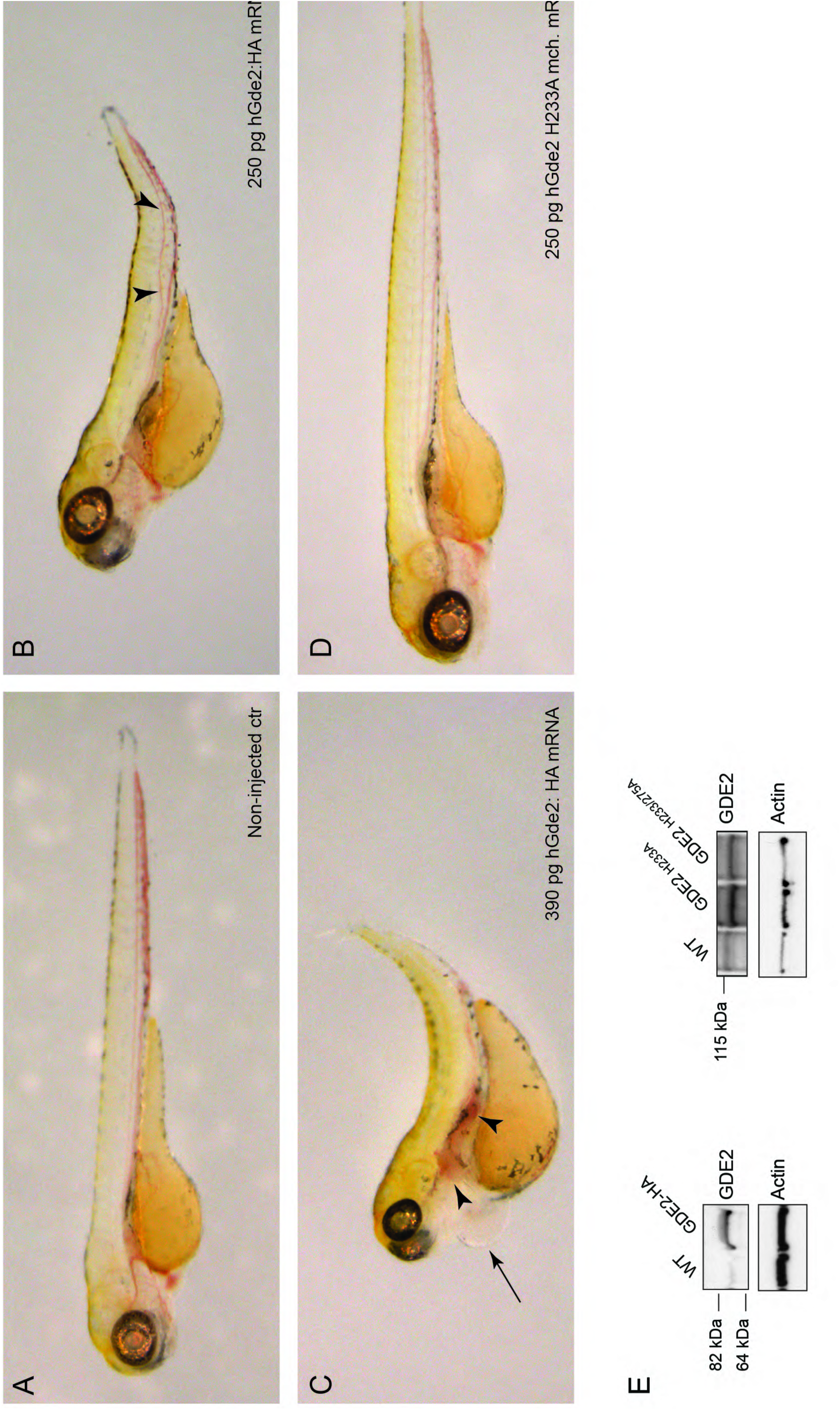
Expression of human GDE2 leads to developmental defects, depending on GDE2 catalytic activity. Phenotypes of larvae at 4 dpf that have been injected at the one-cell stage with mRNAs encoding human GDE2:HA (B, C), or catalytically-dead hGDE2 ^H233A^ mRNAs (D). Lateral views, anterior to the left. **(A)** Non-injected larva at 4 dpf. **(B)** Larva injected with 250 pg of wt hGDE2:HA mRNA. The body axis is shorter with an upward curvature, and defects in the organization of posterior vasculature can be observed (arrowheads). **(C)** Larva injected with 390 pg of hGDE2:HA. The defects are stronger, the body axis is severely shortened, there is blood accumulation and disruption of the posterior vasculature (arrowheads), as well as heart oedemas (arrow). **(D)** Larva injected with 250 pg of the catalytically-dead hGDE2^H233A^; mcherry mRNA. No obvious morphological abnormalities are observed. **(E)** Western-blot analysis of zebrafish at 4 dpf, expressing HA-tagged wt human GDE2, or mCherry tagged GDE2 catalytic dead mutant, GDE2^H233A^. Actin is used as a loading control.

## Discussion

In this study, we have identified a previously unknown role for glycerophosphodiesterase Gde2 (Gdpd5) in non-neuronal cell types, particularly the pancreas. Here, we focused on the possible role of Gde2 during pancreas development in zebrafish and show that Gdpd5 knockdown leads to defects in the differentiation of specific endocrine and exocrine progenitors.

Gdpd5-knockdown leads to dramatic reduction in pancreatic markers trypsin (which marks exocrine cells) and insulin. Other markers of the endocrine pancreas, such as the transcription factor *pdx1* were not affected upon Gdpd5 depletion, indicating that Gdpd5 specifically affects insulin expression rather than the development of the entire endocrine pancreas. Our results therefore strongly suggest that Gdpd5 affects terminal differentiation of insulin-producing beta-cells. With regard to the exocrine pancreas, expression of *ptf1a was* not affected by Gdpd5a knockdown. Trypsin expression, however, was strongly reduced. Interestingly, GDPD5 is also expressed in the human pancreas (Uhlen et al., 2015) (Lang et al., 2008). However, a role for Gdpd5 in pancreas differentiation has not previously been reported. It is of note that ectopic activation of Notch1 signaling inhibits acinar cell differentiation but not initial commitment to the exocrine lineage (Esni et al., 2004). We therefore suggest that *gdpd5a* knockdown affects acinar cell differentiation likely through upregulation of Notch signaling. Consistent with this, *islet1,* a marker of early pancreas development, is not affected upon *gdpd5* knockdown, indicating that Gdpd5 affects differentiation of specific subtypes of pancreatic cells.

Gdpd5a knockdown in zebrafish induces developmental defects, including a short and curved body axis and heart defects. We observed reduced *gdpd5a* expression in the heart of *gdpd5a* morphants, which may lead to increased Notch1 activity in the heart and account for the abnormalities. In chicken embryos and in mice, Gdpd5 (GDE2) downregulates Notch signaling in motor neuron progenitors. It was previously shown that GDE2 promotes neuroblastoma cell differentiation (Matas-Rico et al., 2016a). In adult mice, GDE2 prevents neurodegeneration by promoting motor neuron survival (Cave et al., 2017). Mechanistically, GDE2 does so by specifically cleaving GPI-anchored Notch ligand regulator RECK as well as glypicans at the plasma membrane (Matas-Rico et al., 2016a) (Cave et al., 2017). In zebrafish, Glypicans are highly conserved and affect various signaling pathways including the Wnt and Bmp signaling axis (Gupta et al., 2013;Strate et al., 2015). Interestingly, Gpc4 depletion in zebrafish results in strongly reduced cardiomyocyte proliferation (Strate et al., 2015). Similarly, perturbations in Notch signaling affect cardiomyocyte proliferation during heart development (High et al., 2008) and regeneration (Zhao et al., 2014). Future studies should reveal if the observed heart malformations after perturbed Gde2 expression are associated with impaired Gpc4 or Notch signaling.

Given that Gdpd5 promotes motor neuron differentiation in chicken embryos and mice (Rao et al., 2005) (Sabharwal et al., 2011), while *gdpd5a* morphants show severe mobility defects, it was unexpected to find *islet1* expressed at relatively normal levels in the brain and spinal cord motor neurons in the morphants. A possible explanation could be that motor neurons are specified normally in the Gdpd5a-depleted zebrafish but undergo impaired terminal differentiation downstream of *islet1.* In addition, protein downregulation achieved by morpholino injections is not complete.

Exogenous expression of human GDE2 mRNA similarly led to developmental abnormalities in zebrafish, namely body axis malformations and vasculature and heart defects, which was strictly dependent on GDE2 catalytic activity. These results highlight the functional conservation between human and zebrafish Gdpd5, and indicate that GDE2 expression levels must be tightly regulated for proper development.

Because of the functional conservation between human and zebrafish Gdpd5, the use of zebrafish is an effective model to elucidate the precise signaling axis of GDE2 in pancreas development with potential therapeutic value to diabetes.

## Materials and Methods

### Zebrafish strains and genotyping methods

Adult zebrafish were maintained at 28**°**C in compliance with the local animal welfare regulations. Their culture was approved by the local animal welfare committee (DEC) of the University of Leiden and all protocols adhered to the international guidelines specified by the EU Animal Protection Directive 2010/63/EU. Embryos were staged according to (Kimmel et al., 1995).

### mRNA and Morpholino injections

For mRNA injections, GDE2 constructs previously described in (Matas-Rico et al., 2016a) were used. Briefly, a GDE2 image clone was used as template for PCR amplification and BamH1 and EcoRV were used to clone human full-length GDE2 into pcDNA3 vector containing a C-terminal mCherry or HA tag. The vectors were linearized with Xbal. Capped mRNA was synthesised using the SP6 mMessage mMachine kit (Ambion). mRNA (200-500 pg) encoding either wt hGde2:HA or hGde2^H233A^ was injected into one-cell stage zebrafish embryos. Translation-blocking morpholino (MO) directed against *gdpd5a* morpholino (5′ GTTTCACCATAGTCAGGCCACAGC 3′) was obtained from Gene-Tools (Oregon, USA). Embryos were injected at the 1-2-cell stage with 4- 8,4 ng of MO.

### Whole-mount *in situ* hybridization

Whole-mount in situ hybridizations were carried out according to standard protocols (Thisse et al., 2008). Riboprobes against *Notch1b, ins, trypsin* have been previously described. Plasmids encoding *pdx1* and *ptfa1* were a kind gift from Dr. Ruben Marin Juez. The probe against *islet1* was generated from cDNA clone MGC: 73031, IMAGE: 4144017 (Source Bioscience Lifesciences, Germany), which was linearized with Notl and transcribed using SP6 RNA Polymerase. To generate a riboprobe against *gdpd5a,* an 870 bp fragment was amplified from zebrafish cDNA using the following primers: Forward: 5′ CAGGTTGTAACTCTGGCGGT 3′, Reverse: 5′ TGGGACGAGGCACTTTCTTC 3′. The fragment was cloned into the PGEMT vector, was linearized with Xhol and was transcribed using SP6 RNA polymerase.

### Western Blot analyses

Approximately 20 larvae/sample were lysed (3 μl per larva) in ice-cold standard RIPA buffer supplemented with proteinase inhibitors. Lysates were dounced for 5 min and homogenized using an insulin syringe, followed by centrifugation at 13.000 rpm for 15 min at 4 °C to pellet nuclei and cell debris. The protein concentration was measured using a standard BCA protein assay kit (Pierce). Lysates containing 4x Bolt LDS sample buffer, supplemented with DTT were boiled for 5 min. 12 - 30 ug was loaded onto 12% Bis-Tris SDS-PAGE precast gel (Nu-Page Invitrogen) and transferred to nitrocellulose membranes. Nonspecific protein binding was blocked using 5% skimmed milk in TBST. Primary antibodies were incubated overnight at 4C followed by 1hr incubation with HRP-conjugated secondary antibodies (DAKO, Glostrup, Denmark) and detection using ECL Westernblot reagent (GE Healthcare). Antibodies used were: home-made rabbit anti-GDE2 (Matas-Rico et al., 2016a), mouse anti-beta-actin (1:10.000, Sigma, #A5441).

## Acknowledgements

We thank Dr. Elisa-Matas Rico (NKI, Amsterdam) for GDE2 studies, Dr. Rubén Marín-Juez, (MPI, Bad Neuheim Germany) for probes, and the animal caretakers at Leiden University for excellent care of the fish. The work was supported by grants from the Dutch Cancer Society (KWF UL 2012-5395) to APGH, and Netherlands Organization for Scientific Research (NWO; TOPGO 700.10.354) to WHM.

